# LOCALIZER: subcellular localization prediction of both plant and effector proteins in the plant cell

**DOI:** 10.1101/092726

**Authors:** Jana Sperschneider, Ann-Maree Catanzariti, Kathleen DeBoer, Benjamin Petre, Donald M. Gardiner, Karam B Singh, Peter N. Dodds, Jennifer M. Taylor

## Abstract

Pathogens are able to deliver effector proteins into plant cells to enable infection. Some effectors have been found to enter subcellular compartments by mimicking host targeting sequences. Although many computational methods exist to predict plant protein subcellular localization, they perform poorly for effectors. We introduce LOCALIZER for predicting plant and effector protein localization to chloroplasts, mitochondria, and nuclei. LOCALIZER shows greater prediction accuracy for chloroplast and mitochondrial targeting compared to other methods for 652 plant proteins. For 108 eukaryotic effectors, LOCALIZER outperforms other methods and predicts a previously unrecognized chloroplast transit peptide for the ToxA effector, which we show translocates into tobacco chloroplasts. Secretome-wide predictions and confocal microscopy reveal that rust fungi might have evolved multiple effectors that target chloroplasts or nuclei. LOCALIZER is the first method for predicting effector localisation in plants and is a valuable tool for prioritizing effector candidates for functional investigations. LOCALIZER is available at http://localizer.csiro.au/.

## Introduction

Plant cells feature subcellular compartments such as mitochondria or chloroplasts which contain distinct suites of proteins related to their specialised biological functions. Plant proteins are translocated from the cytosol into specific organelles by means of N-terminal transit peptides in the case of chloroplasts and mitochondria^1,2^, or nuclear localization signals (NLSs) in the case of nuclei. Transit peptides rarely share sequence conservation and vary in length^1,3^.

Eukaryotic filamentous plant pathogens deliver cytoplasmic effectors into host tissues to subvert plant functions to their advantage^4^. Whilst there are many bacterial effectors targeting specific compartments, the extent to which eukaryotic effectors enter plant organelles is less understood^5–11^. Recently, several rust effector candidates were shown to mimic transit peptides to translocate into chloroplasts^12^. Determining effector subcellular localization in plant cells provides important clues about their virulence function. However, experimental methods are labourintensive, prone to artefacts, and not appropriate for high-throughput screening of the large effector repertoires predicted in fungi and oomycetes. No method for effector subcellular localization prediction is available thus far to prioritize effector candidates for experimental investigations.

In principle, a plant-trained classifier should work on both host and pathogen, as effectors might exploit the plant machinery to enter organelles. However, signal peptides, pro-domains and the rapid evolution of effectors pose challenges (Fig. 1). Firstly, if effectors carry transit peptides these can be separated from N-terminal signal peptides by a pro-domain of varying length^13^. This poses a challenge to plant predictors such as TargetP^14^ or ChloroP^15^, which analyse the N-terminal amino acid composition and assume that potential transit peptides start at the first residue after the signal peptide. For example, the ToxA effector localizes to chloroplasts, interacts with the chloroplast-localized protein ToxABP1^16^ and contains a signal peptide followed by a pro-domain that are cleaved during secretion^17^. Thus, if ToxA carries a transit peptide it is likely to occur after the pro-domain. The second challenge lies in the lack of sequence homology between effector proteins and non-effector proteins. Methods such as WoLF PSORT^18^ include homology-based information from proteins with experimentally validated subcellular localizations. However, effectors rarely share sequence similarity with other proteins^19^.

**Fig. 1:**
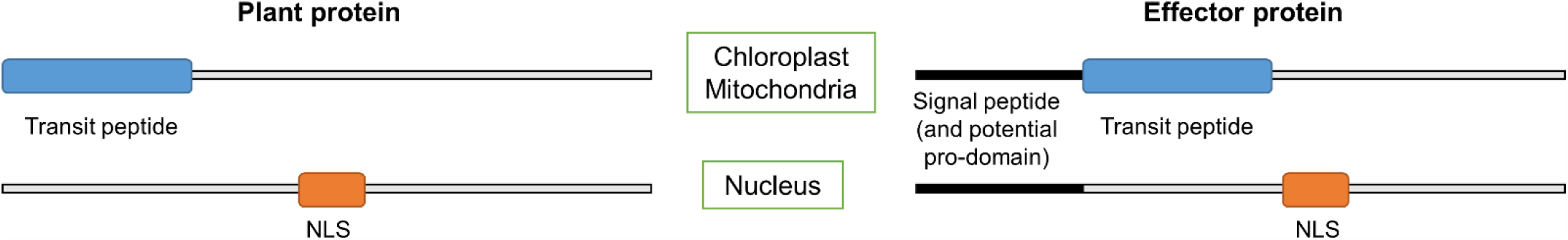
Plant proteins can carry transit peptides at their N-terminus that guide them to chloroplast or mitochondria, or nucleus localization signals (NLSs) that guide them to the nucleus. Effector proteins can also target plant chloroplast, mitochondria, or nuclei through mimicry of transit peptides or NLSs occurring after their signal peptides. Some effectors might also have pro-domains after the signal peptide that are cleaved off during or after secretion.

To address these challenges, we introduce a machine learning prediction tool called LOCALIZER, which can be run in two modes:

1. Plant mode: LOCALIZER predicts chloroplast/mitochondrial transit peptides and/or NLSs in plant proteins.
2. Effector mode: LOCALIZER uses a sliding window approach to predict chloroplast/mitochondrial transit peptides and/or NLSs in effectors.

In both modes, LOCALIZER can predict if a plant or effector protein can localize to multiple compartments. This is important because dual targeting of plant proteins is common^20–22^, but only WoLF PSORT and YLoc+ are capable of predicting dual localization^23^.

## Results

### LOCALIZER: a method for predicting plant and effector protein targeting to chloroplasts, mitochondria or nuclei

To predict if a plant or an effector protein localizes to chloroplasts or mitochondria, we applied a machine learning approach trained on plant proteins with experimentally verified localization data. LOCALIZER uses two classifiers trained to recognize chloroplast or mitochondrial transit peptides and a NLS search for predicting nucleus localization (Supplementary Table S1).

LOCALIZER has been trained on transit peptides and therefore, a window of sequence harbouring a potential transit peptide should be presented to the classifier. To search for transit peptides in plant proteins, sequence windows of varying lengths starting at the first position in the sequence were used. For each of the windows, the chloroplast and mitochondrial classifiers were called. If only one of the classifiers returns predicted transit peptides, the transit peptide with the highest probability is returned as the result (Fig. 2a). If both classifiers return predicted transit peptides, the one with the highest probability is returned as the main localization, e.g. chloroplast, and the weaker classification from the other classifier is reported as a possible dual-localization, e.g. chloroplast and possible mitochondrial (Fig. 2b).

**Fig. 2:**
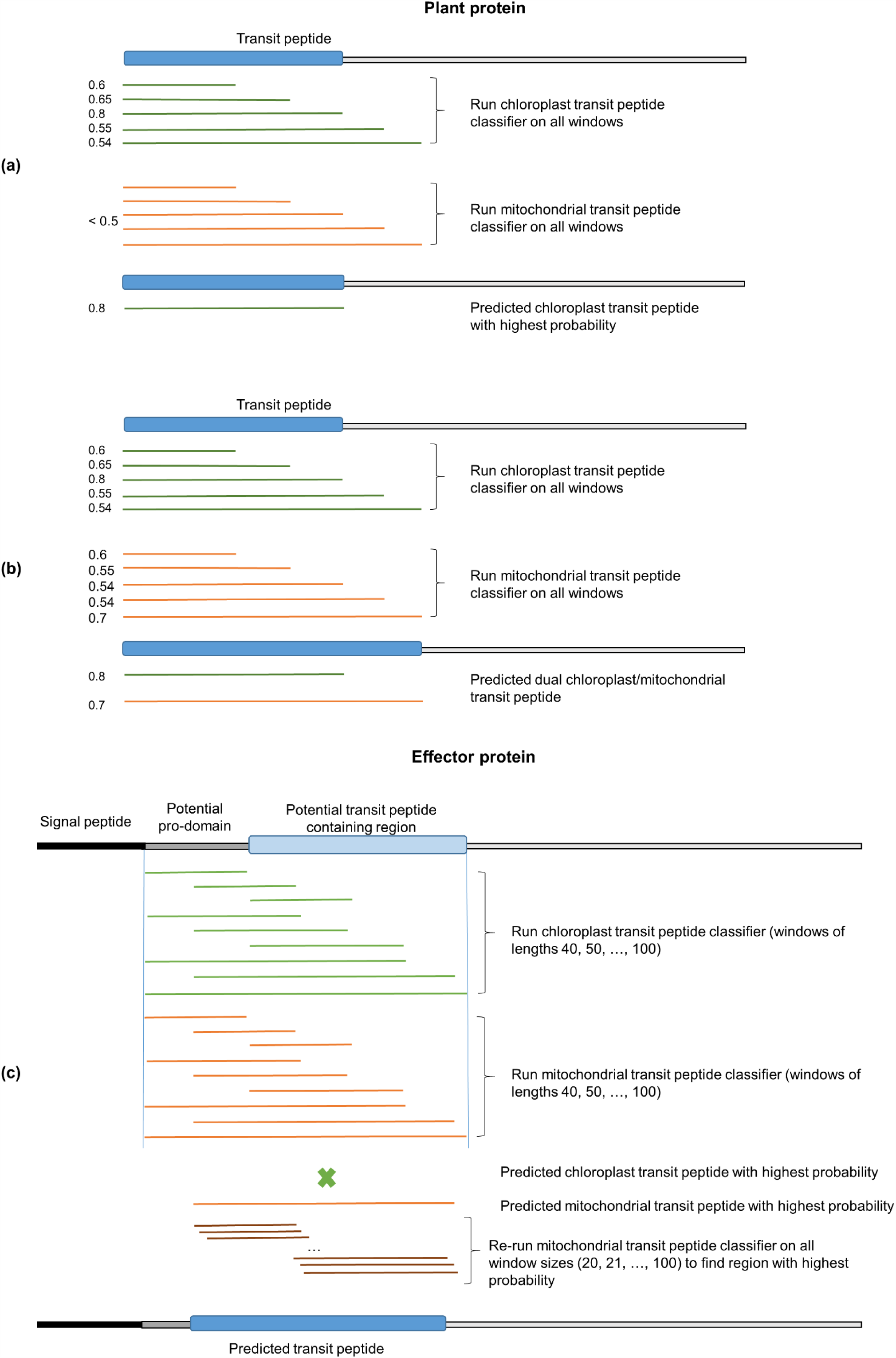
**(a)** An example of a predicted chloroplast-targeting plant protein with probability 0.8. Only the chloroplast classifier returns predicted transit peptides, whereas the mitochondrial classifier does not return positive predictions (probabilities < 0.5). **(b)** An example of a predicted dual-targeting plant protein with a predicted chloroplast transit peptide (probability 0.8) and a possible mitochondrial transit peptide (probability 0.7). **(c)** An example of the sliding window approach for effector proteins. Sliding windows of varying lengths are moved along the mature effector sequences to find potential transit peptide start positions. Both chloroplast and mitochondrial transit peptides are predicted for each window using the two classifier. In this example, the effector has a predicted mitochondrial transit peptide, but no predicted chloroplast transit peptide (indicated by a cross).

Unlike in plant proteins, where the transit peptides starts at the N-terminus, the transit peptide start in effectors is more variable due to signal peptide lengths and the potential presence of pro-domains. To address this, LOCALIZER uses a sliding window approach to scan for the start of potential transit peptides (Fig. 2c). In the following, we first benchmark LOCALIZER on plant proteins and then demonstrate that LOCALIZER also accurately predicts effector localization in the plant cell.

### LOCALIZER has the highest accuracy for predicting chloroplast and mitochondrial localization in plant proteins

Due to the relatively small number of plant proteins with experimentally validated transit peptides in the UniProt database, we could not use parts of our training set as independent test sets. Therefore, we took the manually curated cropPAL database which features the subcellular localization of rice, wheat, maize and barley proteins^24^. We used only those proteins that are supported by GFP studies and localize to one subcellular compartment, resulting in a test set of 530 sequences. The overlap with our UniProt training set is minimal, out of 100 cropPAL proteins that are targeted to plastids only four are part of our chloroplast training set and out of 61 cropPAL proteins that are targeted to mitochondria only four are part of our mitochondria training set. The test set is of high quality due to the manual curation and is also very valuable because it exclusively features crop species, which are host species for the most intensively studied plant pathogens. As a second test set, we used plant proteins with experimentally validated localization that were entered into UniProt after the training of LOCALIZER. This independent test set contains 122 plant proteins.

On the test set of 652 plant proteins, LOCALIZER achieves the highest MCC of 0.71 and highest accuracy of 91.4% for chloroplast proteins and the highest MCC of 0.54 and highest accuracy of 91.7% for mitochondrial proteins compared to other programs (Table 1). However, the homology- and annotation-based methods YLoc and WoLF PSORT achieve higher accuracy of 79.8% and BaCelLo achieves higher MCC of 0.42 on the nuclear localized protein set than LOCALIZER (accuracy 73%, MCC 0.4). The protein property based predictors BaCelLo, NLStradamus and PredictNLS^25^ have lower accuracies than LOCALIZER. PredictNLS in particular correctly identifies only a small proportion (8.9%) as nuclear-localized, suggesting that its default collection of NLSs is incomplete for plants. Note that for LOCALIZER’s evaluation we only considered proteins predicted to localize to the nucleus exclusively in this benchmark. When counting all proteins that are predicted to carry a NLS as nuclear-localized regardless of an additional transit peptide prediction, LOCALIZER’s sensitivity increases, however specificity also decreases. Taken together, this evaluation might suggest that homology-based methods are advantageous for nuclear localization prediction in plants, as a NLS motif search in plant proteomes has a higher false positive rate. Homology-based methods could be combined with a NLS motif search used by LOCALIZER for increased sensitivity in nuclear localization prediction for plant proteins.

**Table 1:** Performance evaluation on the combined test set of cropPAL crop plant proteins and UniProt plant proteins. MCC stands for Matthews Correlation Coefficient and PPV for positive predictive value.

|  |  | Sensitivity | Specificity | PPV | MCC | Accuracy |
| --- | --- | --- | --- | --- | --- | --- |
| **Chloroplast (120 proteins)** |  |  |  |  |  |  |
|  | LOCALIZER | 72.5% | 95.7% | 79.1% | **0.71** | **91.4%** |
|  | TargetP | 75% | 92.9% | 70.3% | 0.66 | 89.6% |
|  | ChloroP | 72.5% | 86.7% | 55.1% | 0.53 | 84% |
|  | WoLF PSORT | **80.8%** | 73.1% | 40.4% | 0.43 | 74.5% |
|  | Predotar | 67.5% | **95.9%** | **78.6%** | 0.67 | 90.6% |
|  | YLoc | 77.5% | 81.2% | 48.2% | 0.5 | 80.5% |
|  | BaCelLo | 75% | 80% | 46.6% | 0.47 | 79.1% |
| **Mitochondria (65 proteins)** |  |  |  |  |  |  |
|  | LOCALIZER | 60% | 95.2% | 58.2% | **0.54** | **91.7%** |
|  | TargetP | **64.6%** | 89.1% | 39.6% | 0.44 | 86.7% |
|  | WoLF PSORT | 21.5% | 95.7% | 35.9% | 0.22 | 88.3% |
|  | Predotar | 60% | 94.4% | 54.2% | 0.52 | 91% |
|  | YLoc | 46.2% | 94% | 46.2% | 0.4 | 89.3% |
|  | BaCelLo | 6.2% | **99.7%** | **66.7%** | 0.18 | 90.3% |
| **Nucleus (225 proteins)** |  |  |  |  |  |  |
|  | LOCALIZER | 60% | 79.9% | 61.1% | 0.4 | 73% |
|  | WoLF PSORT | 55.1% | 92.7% | **80%** | 0.53 | **79.8%** |
|  | YLoc | 61.8% | 89.2% | 75.1% | **0.54** | **79.8%** |
|  | BaCelLo | **82.2%** | 62.1% | 53.3% | 0.42 | 69% |
|  | NLStradamus | 32.9% | 86.9% | 56.9% | 0.24 | 68.3% |
|  | PredictNLS | 8.9% | **98.6%** | 76.9% | 0.18 | 67.6% |

Of the 120 chloroplast-localized proteins, our classifier identifies 87 as carrying a chloroplast transit peptide. Of these 87 proteins, 8 are predicted to also carry a potential mitochondrial transit peptide, 33 are predicted to carry an additional NLS and five are predicted to carry a potential mitochondrial transit peptide as well as an NLS. We used the cropPAL database to build a set of plant proteins that have been shown to localize to two or more compartments based on GFP studies. Out of 34 proteins that have been experimentally shown to localize to both chloroplasts and mitochondria, LOCALIZER correctly predicts dual localization for 13 (38.2%). In contrast, YLoc+ predicts only 6 as dual-targeted to chloroplasts and mitochondria, whereas WoLF PSORT does not assign any of the 34 proteins to the dual compartment chloroplast-mitochondria. Other examples from the literature are the dual-localized maize CFM6 protein^26^, for which LOCALIZER correctly predicts dual localization to nucleus and mitochondria and the WHIRLY1 protein^27^, for which LOCALIZER correctly predicts dual localization to nucleus and chloroplasts. Overall, the independent validation on the test set shows that LOCALIZER outperforms other methods for predicting chloroplast and mitochondrial transit peptides in plant proteins as well as dual-targeting of plant proteins, whereas YLoc and WoLF PSORT are the most accurate tools for plant nuclear localized proteins.

### LOCALIZER in effector mode: high accuracy on fungal and oomycete effector localization

We used data sets from the literature to compare the performance of LOCALIZER to other methods on 108 GFP-tagged fungal and oomycete effectors/effector candidates (Supplementary Table S2, results given in Supplementary File 2, bacterial effector prediction described in Supplementary File 1). Out of the 108 effectors, seven effectors have been shown to localize to chloroplasts and 51 effectors have been shown to localize to the plant nucleus. To the best of our knowledge, no localization to mitochondria exclusively has been shown for eukaryotic effectors from plant pathogens. We tested LOCALIZER in effector mode and compared it to the performance of plant subcellular localization methods applied to mature effector sequences. Overall, LOCALIZER had the best performance when predicting chloroplast- or nucleus-localized effectors, whereas plant subcellular localization methods showed low sensitivity, MCCs close to zero and high false positive rates (Table 2) and are therefore not suitable for predicting effector localization. Notably, the chloroplast-targeting effector set demonstrates that plant subcellular localization prediction methods should not be used for predicting the chloroplast-targeting of mature effector sequences due to poor performance. LOCALIZER predicts chloroplast transit peptides for 5 out of 7 effectors (Table 3), whereas other methods rarely predict chloroplast localization in these effectors and feature higher false positive rates of 5% to 21.8% (Table 2).

**Table 2:** Evaluation of effector localization using LOCALIZER and other plant-trained subcellular localization methods.

| Chloroplast effectors |  | Sensitivity | Specificity | PPV | MCC | Accuracy | False positive rate |
| --- | --- | --- | --- | --- | --- | --- | --- |
|  | LOCALIZER | **85.7%** | **96%** | **60%** | **0.69** | **95.4%** | **4%** |
|  | TargetP | 0% | 83.2% | 0% | -0.11 | 77.8% | 16.8% |
|  | ChloroP | 14.3% | 92.1% | 11.1% | 0.06 | 87% | 7.9% |
|  | WoLF PSORT | 42.9% | 78.2% | 12% | 0.12 | 75.9% | 21.8% |
|  | BaCelLo | 0% | 88.1% | 0% | -0.09 | 82.4% | 11.9% |
|  | YLoc | 14.3% | 92.1% | 11.1 | 0.06 | 87% | 7.9% |
|  | Predotar | 0% | 95% | 0% | -0.06 | 88.9% | 5% |
| Nucleus effectors |  |  |  |  |  |  |  |
|  | LOCALIZER | 51% | 87.7% | 78.8% | **0.42** | **70.4%** | 12.3% |
|  | WoLF PSORT | 47.1% | 75% | 63.2% | 0.22 | 61.1% | 25% |
|  | BaCelLo | 52.9% | 62.5% | 56.3% | 0.15 | 57.4% | 37.5% |
|  | YLoc | **58.8%** | 75% | 68.2% | 0.33 | 66.7% | 25% |
|  | NLStradamus | 23.7% | 94.7% | 82.4% | 0.3 | 63% | 5.3% |
|  | PredictNLS | 11.8% | **100%** | **100%** | 0.15 | 54.6% | **0%** |

**Table 3:** Test set of effectors and effector candidates that have been experimentally shown to localize to the plant chloroplast and the predictions by several classifiers. All methods except LOCALIZER were supplied with the mature effector sequences.

| Effector | CTP1 | CTP2 | CTP3 | MLP124111 | PST03196 | PST18220 | ToxA |
| --- | --- | --- | --- | --- | --- | --- | --- |
| Reference | Petre *et al.* (2015b), Petre *et al.* (2015a) |  |  |  | Petre *et al.* (2016) |  | Manning and Ciuffetti (2005) |
| Species | *Melampsora larici-populina* |  | *M. lini* | *M larici-populina* | *Puccinia striiformis* f. sp. *tritici* |  | *Pyrenophora tritici-repentis* |
| Comment | Also in mitochondria | - | - | Also in cytosolic aggregates |  | Also in nucleus | Also in cytoplasm |
| Subcellular localization prediction |  |  |  |  |  |  |  |
| LOCALIZER | Chloroplast | Chloroplast | Chloroplast | - | Chloroplast, Mitochondria, Nucleus | - | Chloroplast |
| ChloroP | Chloroplast | - | - | - | - | - | - |
| TargetP | Mitochondria | Mitochondria | - | Mitochondria | - | - | - |
| WoLF PSORT | Chloroplast | Chloroplast | Chloroplast | Mitochondria | Cytosol | Nucleus | Cytoskeleton |
| YLOC | Secreted | Secreted | Secreted | Secreted | **Chloroplast** | Nucleus | Cytoplasm |
| Predotar | - | - | - | - | - | - | - |
| BaCelLo | Secreted | Secreted | Secreted | Secreted | Nucleus | Nucleus | Nucleus |

We then tested LOCALIZER on the set of 51 effectors that have been experimentally shown to localize to the plant nucleus. On this set, LOCALIZER had the highest MCC of 0.42 and highest accuracy of 70.4% (Table 2). Despite their superior performance on plant proteins, BaCelLo, WoLF PSORT and YLoc have lower MCCs and accuracies for nucleus-localized effectors. In particular, the majority of *Hyaloperonospora arabidopsis* RxLR (HaRxLR) effector candidates in Caillaud, et al.^6^ were experimentally found to localize to the nucleus, whereas none have been found to enter chloroplasts and mitochondria. LOCALIZER predicts NLSs in 7 out of 13 HaRxLR effectors (53.8%) that exclusively localize to the nucleus, but only in 3 out of 15 HaRxLR effectors (20%) that have been shown to localize to the nucleus and cytoplasm, which might indicate that their localization to nuclei is inconclusive. For the set of 45 HaRxLR effector candidates, LOCALIZER returns seven false positive predictions of chloroplast or mitochondrial targeting (15.6%). Interestingly, when providing LOCALIZER with HaRxLR mature effector sequences that are cleaved after the leucine in the RxLR motif, this changes to two false positives (4.4%). Our negative set also includes 33 candidate rust effectors that show no specific localization to mitochondria, chloroplast or nuclei in experiments^12,28,29^. For these 33 effector candidates, LOCALIZER only predicts three as nuclear-localized and one as chloroplast-localized, whereas ChloroP predicts five as chloroplast-localized and TargetP returns 11 false positives. Overall, these results show that plant subcellular localization methods are not suitable for predicting effector localization and that LOCALIZER greatly improves prediction accuracy for effectors targeting plant chloroplasts and nuclei.

### LOCALIZER predicts a chloroplast transit peptide for ToxA and suggests that many RxLR effectors target nuclei

We collected a set of experimentally validated 69 fungal effectors and 51 oomycete effectors and ran LOCALIZER to predict their subcellular localization. For the 69 fungal effectors, only three are predicted to carry a chloroplast transit peptide, i.e. Six6 (*Fusarium oxysporum* f. sp. *lycopersici*), Ave1 (*Verticillium dahliae*) and ToxA (*Pyrenophora tritici-repentis* and *Parastagonospora nodorum*). Both Six6 and Ave1 are likely false positives, Six6-GFP has been shown to localize to the cytoplasm and cell nucleus in tobacco^30^ and Ave1 is recognized on the cell surface. Interestingly, Ave1 has been suggested to be horizontally transferred from plants^31^. Only the Avra10 effector from *Blumeria graminis* f.sp. *hordei* is predicted to target mitochondria and also has a predicted NLS. Nine effectors are predicted to carry a NLS (RTP1, Avr4, Avr2, Six1, Bas107, Avr-Pik, UhAvr1, Pit2, SP7; 13% of effectors). Of these, RTP1, SP7 and Bas107 have thus far been shown to localize to the plant nucleus^32–34^. We could not predict nuclear localization for the fungal effectors MISSP7^35^, See1^36^ and Six3^37^, suggesting that these might not rely on NLSs to enter plant nuclei.

ToxA has been implicated to localize to chloroplasts, but the presence of a transit peptide has thus far not been confirmed using existing prediction methods^16,38^. However, LOCALIZER predicts a chloroplast transit peptide at position 62 to 130 with probability 0.885 for the *Parastagonospora nodorum* ToxA. Mapping the chloroplast transit peptide predicted by LOCALIZER onto the ToxA structure reveals that it is predicted to start immediately after the pro-domain (Fig. 3). The pro-domain of ToxA has been suggested to finish at amino acid 60 and to be important for folding, but not necessary for toxic activity^39^. We found that ToxA_62-132_-GFP accumulates in *N. benthamiana* chloroplasts (Fig. 3). A shorter version of the predicted transit peptide (ToxA_62-93_-GFP), where amino acid position 93 is the start of the first beta sheet in the three-dimensional structure^40^ did not localize to tobacco chloroplasts (Fig.3). Furthermore, the full-length ToxA protein (with and without the pro-domain) did not localize to tobacco chloroplasts (data not shown). Taken together, this suggests that the N-terminal region of ToxA (ToxA_62-132_) has the ability to enter chloroplasts, however it likely needs to be unstructured in order to do so, which might be achieved through the pro-domain or post-translational modifications.

**Fig. 3:**
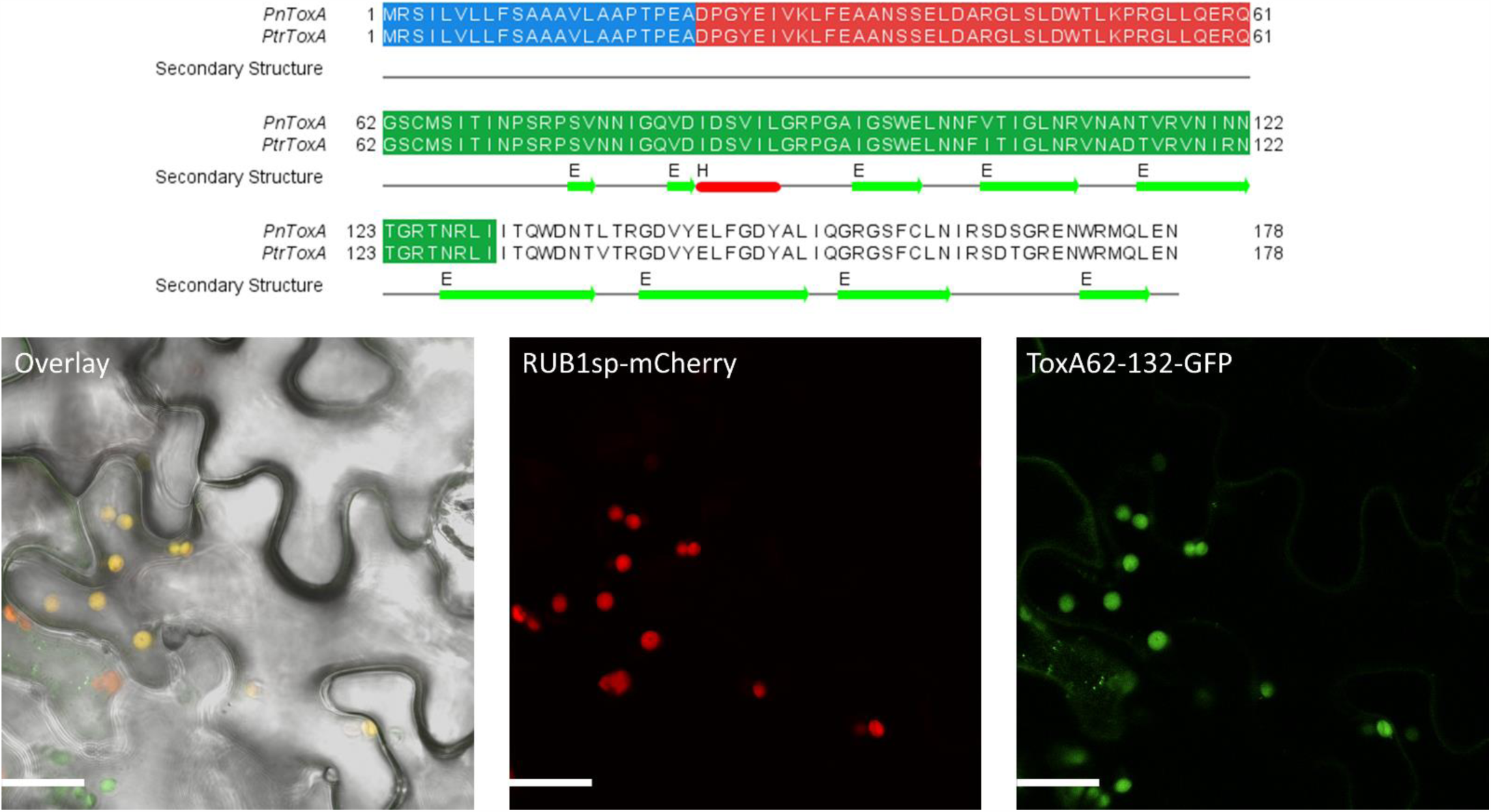
The sequence that was tested for chloroplast localization for PnToxA (*Parastagonospora nodorum*) is shown in green in the alignment. The signal peptide and pro-domain are shown in blue and red, respectively. The secondary structure from PtrToxA (PDB entry 1zle) is also shown. Confocal images demonstrating the chloroplast localisation of ToxA_62-132_-GFP when transiently expressed in the pavement cells of *N. benthamiania* leaves. The left panel displays the transmitted light image overlayed with the ToxA_62-132_-GFP and RUB1sp-mCherry fluorescence. The middle panel displays the RUB1sp-mCherry fluorescence (chloroplast marker) whilst the right panel displays the ToxA_62-132_-GFP fluorescence. Scale bar = 20 μm

For predictions on the oomycete effector set we used RxLR mature effector sequences cleaved after the leucine in the RxLR motif. Only the *Phytophthora sojae* Pslsc1 effector is predicted as carrying a chloroplast transit peptide, which may be consistent with its function in suppressing salicylate-mediated immunity by degrading isochorismate (produced in the chloroplast)^41^. The RxLR effector Avh241 is predicted to target mitochondria but has been shown to localize to plasma membranes^42^. A large proportion of oomycete effectors are predicted to carry NLSs (ATR13, Avrblb2, CRN1, CRN2, CRN8, CRN15, CRN16, CRN63, CRN115, Avh18a1, PiAvr2, PiAvrVnt1, PsAvr3b, PsAvr4/6; 27.5% of effectors). Several Crinkler (CRN) effector proteins have been shown to localize to the plant nucleus and to require nuclear accumulation to induce plant cell death^10^. LOCALIZER predicts all seven Crinkler effectors as nuclear-localized, whereas YLoc predicts three, WoLF PSORT and BaCelLo predict two and NLStradamus and PredictNLS predict only one as nuclear-localized. On a set of 358 RxLR effector candidates determined using the Hidden Markov model^43^ and cleaved after the leucine in the RxLR motif, LOCALIZER predicts 0.9% predicted as chloroplast-targeting, 6.1% predicted as mitochondrial-targeting and 22.3% predicted as nucleus-targeting. This suggests that oomycete effectors may not target chloroplasts, but given that a large proportion carry predicted NLSs, may preferably target the plant nucleus.

### Localization predictions of effector candidates reveals expanded chloroplast- and nuclear-targeting in rust pathogens

To further investigate the extent to which fungal effectors are targeted towards specific plant compartments, we first predicted secretomes from 61 fungal species including pathogens and saprophytes using a sensitive approach tailored to effector finding described in Sperschneider, et al.^44^. Importantly, the presence of a predicted signal peptide does not necessarily guarantee the full secretion of a protein from the pathogen. Firstly, prediction tools can return false positives. On eukaryotic data, SignalP 3 (which is used here for secretome prediction) has been estimated to have sensitivity of 98.8% with a false positive rate of 0.8% to 11.7% depending on the presence of transmembrane domains (http://www.cbs.dtu.dk/services/SignalP/performance.php). Secondly, proteins with a signal peptide can have retention signals such as the consensus sequences KDEL/HDEL that can keep them in the ER or Golgi^2^ or proteins with a signal peptide can be anchored to the pathogen cell wall. Therefore, we also ran EffectorP^45^ on the secretomes to limit the subcellular localization predictions to likely effector candidates.

Saprophytes are unlikely to produce effectors targeted to plant subcellular compartments, and indeed only very low numbers of predicted effectors were predicted to show chloroplast (1.24%), mitochondrion (0.73%) or nuclear (1.78%) localisation from these fungi (Table 4, Supplementary Figs. S1-S3). The proportions of predicted chloroplast, mitochondria and nuclear targeted proteins were slightly higher amongst effector candidates predicted from pathogen/symbiont secretomes, with a total of 2.18%, 1.03% and 2.62% respectively. Interestingly, the haustoria-forming pathogens (rusts and powdery mildews) showed even higher proportions of predicted nuclear-targeted effectors (˜4.5%). The highest proportions (> 5%) of effector candidates that were also predicted to target nuclei were found in *Puccinia graminis* f. sp. *tritici* (5.61%), *P. triticina* (5.51%), *Magnaporthe oryzae* (5.46%) and *Blumeria graminis f. sp. hordei* (5.1%). Likewise, the rust pathogens had a higher proportion of predicted chloroplast-targeting effectors (3.35%), while this was not observed for *Blumeria* species (1.03%), which infect only cereal epidermal cells deficient in chloroplast. The apoplastic pathogen *Cladosporium fulvum* showed only low levels of targeted effector candidates similar to the observations for saprophytic fungi and the animal pathogen *Batrachochytrium dendrobatidis* also has a low percentage of predicted chloroplast-targeting effectors of 1.33% (Table 4).

**Table 4:** Percentages of secreted proteins that are EffectorP effector candidates and are also predicted to target chloroplast, mitochondria or nuclei by LOCALIZER across groups of fungal genomes.

|  | Number of genomes | Average % of secreted proteins that are EffectorP effector candidates and are also predicted to target |  |  |
| --- | --- | --- | --- | --- |
|  |  | Chloroplast | Mitochondria | Nucleus |
| **Plant pathogens/symbionts** | 36 | 2.18% | 1.03% | 2.62% |
| **Saprophytes** | 19 | 1.24% | 0.73% | 1.78% |
| **Rust pathogens** | 5 | **3.35%** | 0.93% | **4.5%** |
| **Blumeria pathogens** | 2 | 1.03% | **1.74%** | 4.37% |
| ***Cladosporium fulvum*** | 1 | 1.19% | 0.37% | 2.2% |
| ***Magnaporthe oryzae*** | 1 | 2.58% | 0.8% | 5.46% |
| ***Batrachochytrium dendrobatidis*** | 1 | 1.33% | 0.48% | 3.03% |

Subcellular localization prediction of effector candidates relies strongly on accurate signal peptide prediction. Some fungal proteins might be retained in the fungal cell despite a predicted signal peptide and localize to fungal mitochondria or nuclei. The use of sensitive signal peptide prediction methods is especially advantageous for effector finding^44^ however it might lead to a higher number of false positives. For example, we found that the *Saccharomyces cerevisiae* NAD-dependent protein deacetylase HST1 is predicted as secreted and LOCALIZER predicts it to have an NLS, but is annotated in UniProt as non-secreted and nuclear. Another example is the *S. cerevisiae* fumarate reductase 2 protein which has been shown to localize to mitochondria^46^, but is predicted to be secreted by SignalP 3.0. In effector mode, LOCALIZER predicts chloroplast localization for the mature protein with probability 0.98 whereas in plant mode, LOCALIZER correctly identifies a mitochondrial transit peptide at the start of the protein with probability 0.992. Therefore, the combination of effector candidate prediction with subcellular localization prediction might be advantageous to arrive at a confident set of likely effectors that can target plant cell compartments.

We selected four secreted proteins from the wheat stem rust fungus *P. graminis* f. sp. *tritici* and tested them for their subcellular localization in tobacco. The four candidates were selected based on their differential expression in haustoria compared to spores^47^, their sequence homology to rust pathogens only and their predicted subcellular localization to chloroplast, mitochondria or nucleus based on LOCALIZER and other tools. We ran SignalP 4.1 to determine the signal peptide cleavage site and ran localization predictions on the mature sequences (Supplementary Table S3). The localization experiments showed that the mature sequences of PGTG_00164 and PGTG_06076 accumulate in chloroplasts, whereas PGTG_13278 and PGTG_15899 localize to nuclei (Fig. 4).

**Fig. 4:**
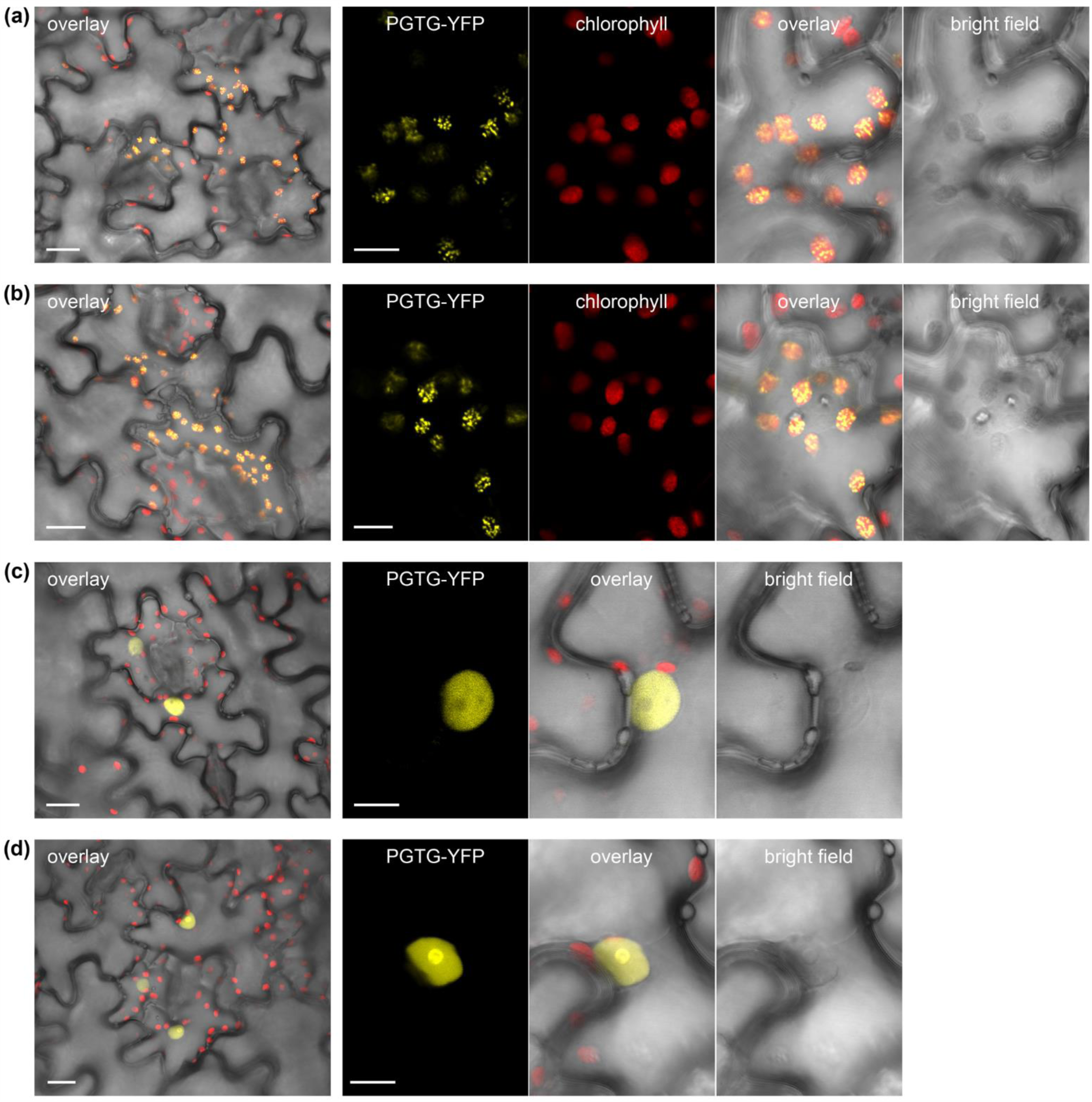
Fluorescence microscopy of YFP-tagged rust effector candidates transiently expressed in tobacco leaves. Confocal images of tobacco cells (*Nicotiana tabacum*) expressing *Puccinia graminis* f. sp. *tritici* (PGTG) effector candidates fused to yellow fluorescent protein (YFP). Cells expressing PGTG_00164-YFP (a) or PGTG_06076-YFP (b) show chloroplast localisation with a punctate pattern suggesting thylakoids or plastoglobules. Cells expressing PGTG_13278-YFP (c) show nuclear localisation with nucleolus exclusion, and those expressing PGTG_15899-YFP (d) show nuclear and nucleolus localisation. Larger panels (left) show transmitted light image of pavement cells overlayed with YFP fluorescence and chlorophyll autofluorescence (overlay); bar = 20 μm. Smaller panels show transmitted light images (bright field) and corresponding fluorescence from PGTP-YFP fusion proteins or chlorophyll (a & b only), and an overlay of all three; bar = 10 μm.

PGTG_00164 is predicted as chloroplast-targeted by YLoc, Predotar and BaCelLo, whereas LOCALIZER detects only a bipartite NLS in the sequence and also in two of its homologs (Supplementary Fig. S5). PGTG_06076 is a larger effector candidate protein of 336 aas and is correctly predicted to target chloroplasts by all tested methods including LOCALIZER. LOCALIZER also predicts chloroplast targeting for the homologs PST130_11623 and PTTG_29922, but not for PGTG_07697 (Supplementary Fig. S6). PGTG_13278 has sequence homology to a large number of proteins in other rust pathogens and is correctly predicted to localize to nuclei by all predictions tools that can predict nucleus localization (Supplementary Table S4). PGTG_15899 localizes to the nucleus and, to a lesser extent, the cytoplasm, but is predicted to target chloroplasts and/or mitochondria by ChloroP, TargetP, WoLF PSORT, YLoc and BaCelLo. LOCALIZER predicts a mitochondrial transit peptide in the mature PGTG_15899 sequence, but also identifies a bipartite NLS (Supplementary Fig. S7).

## Discussion

Plants face attack by pathogens that secrete effector proteins into plant cells to subvert plant defences or manipulate physiology by targeting host proteins. These effectors would be expected to co-localize in subcellular compartments with their respective host targets. For example, plant chloroplasts have emerged as a target of microbial effector proteins^48,49^ and bacterial effectors have been found to directly target mitochondria^50^. The plant nucleus is another prime target for pathogen effectors both from bacteria and eukaryotes^10,11,51^. Furthermore, chloroplast-nucleus communication via stromules has been suggested to play a role in the plant immune response^52^. Accurate predictions of subcellular localization for both plant and pathogen proteins is therefore essential for understanding pathogen-targeted host components and compartments. Many computational subcellular localization prediction tools have been developed for plant proteins, however no dedicated methods are available for predicting effector localization in the plant cell and we found that plant subcellular localization methods on mature effector sequences give unreliable results. Thus, we developed LOCALIZER as the first computational prediction method capable of accurately predicting both the localization of pathogen effector proteins as well as plant proteins to chloroplasts, mitochondria or nuclei.

For nuclear localization prediction, LOCALIZER uses a simple search for eukaryotic NLSs. On the set of plant proteins, LOCALIZER is outperformed by YLoc and WoLF PSORT that include homology-based information due to the low specificity of the NLS search. However, on secreted effector proteins a NLS search employed by LOCALIZER shows the highest accuracy. For effector proteins in particular, using homology to known nuclear-localized proteins can be a misleading strategy as gene families might have members that have evolved different functions or subcellular localization. For example, the rust effector RTP1 from *Uromyces fabae* (Uf-RTP1) has been shown to localize to the plant nucleus, whereas its homolog from *U. striatus* (Us-RTP1) has been found only in the host cytoplasm and is barely visible in nuclei^32^. LOCALIZER predicts nuclear localization for Uf-RTP1, but not for Us-RTP1, whereas YLoc predicts nuclear localization for Us-RTP1. Furthermore, effector proteins rarely share significant similarity to know proteins and thus, homology-based methods underperform on the set of nuclear-localized effectors whereas LOCALIZER delivers the highest accuracy.

On sets of experimentally validated fungal and oomycete effectors, we found that the mimicry of chloroplast and mitochondrial transit peptides seems rare. However, the ability to target chloroplasts can be significant for effector function, as exemplified by the ToxA effector and its interaction with a chloroplast-localized plant protein as well as light-dependency^16,38^. LOCALIZER is the first method which predicts a previously undetected chloroplast transit peptide in the ToxA effector, and we demonstrate that the predicted transit peptide indeed has the ability to enter tobacco chloroplasts. Many experimentally validated effectors, especially from oomycetes, were predicted to target host nuclei. When scanning predicted fungal secretomes for subcellular localization, we found that rust pathogens, *Blumeria* pathogens and *Magnaporthe oryzae* seem to be enriched for predicted nucleus-targeting effector candidates relative to other fungi. Rust pathogens also showed an enrichment for predicted chloroplast-targeting effector candidates relative to other fungi. Using confocal microscopy and transient expression, we confirmed chloroplast-targeting and nucleus-targeting for four rust effector candidates.

We have demonstrated that LOCALIZER will facilitate both functional plant protein and effector studies and improve our understanding of plant-pathogen interactions. Future developments of LOCALIZER will include the capability to predict the localization to other compartments in the plant cell such as the apoplast, peroxisome, cytoplasm and plasma membrane.

## Methods

### Training set selection and support vector machine training

The UniProt data base was accessed and plant proteins (taxonomy:”Viridiplantae [33090]”) were retrieved that localize to the following compartments supported by experimental evidence: chloroplast (”Plastid [SL-0209]”, “Chloroplast [SL-0209]”); mitochondria (”Mitochondrion [SL-0173]”); nucleus (”Nucleus [SL-0191]”); cytoplasm (”Cytoplasm [SL-0086]”); membranes (”membrane”); secreted (”Secreted [SL-0243]”). Sequences that were shorter than or equal to 50 amino acids (aas) and those that did not start with an ‘M’ were removed. From the set of 1,600 chloroplast localized proteins, 1,279 annotated transit peptides were extracted and saved as a FASTA file. The average length of transit peptides was 53 (min:15 max:113). After homology reduction by excluding transit peptides that share sequence similarity with another one at E-value < 0.00001 using phmmer^53^, 639 transit peptides remained in the training set. From the set of 326 mitochondrial localized proteins, 201 annotated transit peptides were extracted and saved as a FASTA file. The average length of transit peptides was 41 (min:12 max:115). After homology reduction, 194 transit peptides remained in the training set.

We also retrieved nuclear-localized eukaryotic proteins from UniProt (taxonomy:“Eukaryota [2759]”, “Nucleus [SL-0191]”, evidence:experimental). The annotated NLSs were extracted and saved as a list of motifs. These were supplemented with a list of regular expressions that describe known NLSs given in Kosugi, et al.^54^ and the bipartite NLS defined as follows: two adjacent basic amino acids (R/K) followed by a spacer region of 8-12 residues followed by at least three basic amino acids (R/K) in the five positions after the spacer region. Additionally, we used NLStradamus, a NLS predictor using Hidden Markov Models which has been trained on yeast sequences^55^. A protein is labelled as containing a NLS if at least one of the two methods are positive: 1) NLS sequence motif search, or 2) NLS prediction by NLStradamus. For each protein, all predicted NLSs are returned and if one motif is contained in another one, the longer NLS is returned as the result. All training sets and the NLS list are available at http://localizer.csiro.au/data.html.

The chloroplast classifier was trained to distinguish chloroplast transit peptides from N-termini of non-chloroplast proteins. The set of 639 chloroplast transit peptides was used as the positive set and the set of homology-reduced non-chloroplast plant proteins was used as the negative set (1,597 proteins). For each protein from the negative set, a random choice of the first *x* amino acids was used, with *x* ranging from 40 to 120. The mitochondria classifier was trained to distinguish mitochondrial transit peptides from N-termini of non-mitochondrial proteins. The set of 194 mitochondrial transit peptides was used as the positive set and the set of homology-reduced non-mitochondrial plant proteins was used as the negative set (1,878 proteins). As the negative set is much larger than the positive set, a randomly chosen subset of 626 proteins was used as the negative set. For each protein from the negative set, a random choice of the first *x* amino acids was used, with *x* ranging from 40 to 120.

The feature vector for each protein consists of 58 features given in Supplementary Table S1. Both classifiers were trained using the SMO support vector machine classifier with the RBF Kernel and the option -M for fitting logistic models using the Weka software (weka.classifiers.functions.SMO)^56^. To optimize the complexity parameter *C* and gamma parameter *G* of the support vector machine, the Weka grid search was used to find the best parameters. For the chloroplast classifier, the best parameters found were *C* = 2.0 and *G* = 1.0 and for the mitochondrial classifier, the best parameters found were *C* = 3.0 and *G* = 0.0625.

### Test sets and performance evaluation

We downloaded the set of crop plant proteins (barley, wheat, rice, maize) from the cropPal database^24^ and chose those that have a subcellular localization of either ‘plastid’ (100 proteins), ‘mitochondrion’ (61 proteins), ‘nucleus’ (165 proteins), ‘peroxisome’ (11 proteins), ‘vacuole’ (18 proteins), ‘plasma membrane’ (84 proteins, ‘endoplasmic reticulum’ (43 proteins) and ‘cytosol’ (48 proteins) determined by GFP-tagging. We only kept those sequences that started with an ‘M’. For the UniProt test set, we downloaded plant proteins (taxonomy:“Viridiplantae [33090]”) that were entered after our training sets were compiled (created:[20160301 TO 20160902]) for several compartments supported by experimental evidence (”Nucleus [SL-0191]”; “Mitochondrion [SL-0173]”, “Chloroplast [SL-0049]”, “Peroxisome [SL-0204]”, “Vacuole”, “Secreted”, “Endoplasmic reticulum”, “Cytoplasm”). We manually removed those entries that localize to multiple compartments, except for the category nucleus for which we also allowed an additional cytoplasmic localization. All plant and effector test sets are available at http://localizer.csiro.au/data.html.

When evaluating performance, the number of true positives (TPs), true negatives (TNs), false positives (FPs) and false negatives (FNs) were used. Sensitivity 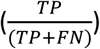 is defined as the proportion of positives that are correctly identified whereas specificity 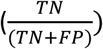 is the proportion of negatives that are correctly identified. Precision (positive predictive value, PPV, 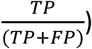 is a measure which captures the proportion of positive predictions that are true. Both accuracy 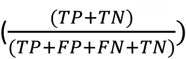 and the Matthews Correlation Coefficient 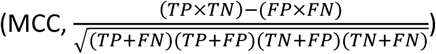 can be used to evaluate the overall performance of a method. The MCC ranges from −1 to 1, with scores of −1 corresponding to predictions in total disagreement with the observations, 0.5 to random predictions and 1 to predictions in perfect agreement with the observations. For our classifier, we count LOCALIZER predictions that are ‘chloroplast’, ‘chloroplast and possible mitochondrial’, ‘chloroplast and nucleus’ and ‘chloroplast & possible mitochondrial and nucleus’ as chloroplast predictions (same strategy for mitochondrial predictions). A protein that carries a predicted transit peptide with an additional predicted NLS might have experimental evidence only for one of those locations due to the technical hurdles of recognizing dual targeting^20^ and should thus not necessarily be counted as a false positive prediction. A protein is counted as a nucleus prediction only if it has the category ‘nucleus’ to avoid assigning a protein to multiple predictions in the evaluation. Many plant subcellular localization methods have been published, however only a small number are available as standalone software or have the option of submitting large batch sequence files to a web server. This makes it prohibitive for researchers to use them routinely for data analysis and thus, our benchmark only includes methods that can be locally installed with ease or have a web server with a batch file submission option (Supplementary Table S2).

### Detection of transit peptides in plant proteins and effectors

In plant mode, windows of varying lengths (40, 50, 60, 70, 80, 90, 100) starting from the first position in the sequence are used to search for chloroplast and mitochondrial transit peptides in plant proteins. In effector mode, sliding windows of varying lengths (40, 50, 60, 70, 80, 90, 100) are moved with step size of one amino acid along the mature effector sequences to find potential transit peptide start positions under the following criteria: 1) to allow for potential pro-domains, the transit peptide can have a start position ranging from residue 1 to 50 in the mature sequence, and 2) at least 40 aas must remain in the C-terminal to allow for the effector domain after the potential transit peptide region. Unless mature effector sequences are provided by the user, LOCALIZER removes the signal peptide by deleting the first 20 aas. In the evaluation, we use the default method of deleting the first 20 aas in the effector sequences unless otherwise specified.

For each of the windows, the chloroplast and mitochondrial classifiers are called and the support vector machine (SVM) probability for a positive classification is returned. To reduce false positive predictions, proteins that have five or less positive predictions in all sliding windows are discarded as weak predictions in effector mode. In both plant and effector mode, a transit peptide prediction is only made if the probability is larger than 0.6. If only one of the classifiers returns predicted transit peptides, the transit peptide with the highest probability is returned as the result. If both classifiers return predicted transit peptides, the one with the highest probability is returned as the main localization and the weaker classification from the other classifier is reported as a possible dual-localization. Up to this point, only certain windows of sizes (40, 50, 60, 70, 80, 90, 100) were considered due to computational runtime and thus only approximate locations of transit peptides are returned. Let *x* be the length of the predicted transit peptide window with highest probability. To identify the location of the predicted transit peptide more accurately, a second round of sliding window predictions (windows of varying lengths from size 20, 21, 22, …, *x*; *x* < 100; step size of one amino acid) are run on the predicted transit peptide window with the highest probability. If one of the refined windows return a transit peptide with a higher probability, this is reported in the final prediction.

If a protein contains more than 10% unknown bases (B, Z, X) in its sequence, it is not used for prediction of localization. If it contains less than 10% unknown bases, these are randomly replaced with the respective amino acid (B replaced with D or N; Z replaced with E or Q, X replaced with any amino acid). In plant and effector mode, only proteins longer than 40 aas will be used for prediction of localization.

### Prediction of fungal pathogen secretomes

The set of secreted proteins was predicted in various fungal genomes (Supplementary File 3) using a pipeline described in Sperschneider, et al.^44^ which assigns a protein as secreted if it is predicted to be secreted by the neural network predictors of SignalP 3 and by TargetP and if it has no predicted transmembrane domain outside the first 60 aas using TMHMM and no predicted transmembrane domain using Phobius^14,57–59^. EffectorP 1.0 was run with default parameters to predict effector candidates from the fungal secretomes^45^.

### Transient *in planta* expression and confocal microscopy of ToxA transit peptide

The ToxA transit peptide was codon optimised for expression in *Nicotiana benthamiania* using the Integrated DNA Technologies (IDT) codon optimisation tool (https://sg.idtdna.com/CodonOpt). The codon optimised ToxA transit peptide was fused to the N-terminus of GFP synthetically as a gBlocks gene fragment (IDT), and was subsequently cloned into the pAGM4723 binary vector, behind the CaMV 35S promoter, using the Golden Gate system^60^. For chloroplast co-localisation, we used the pCMU-PLAr plasmid containing the *AtUBQ10* promoter: RUB1sp-mCherry expression cassette, as described in Ivanov and Harrison^61^. Localisation experiments were undertaken essentially as described in Sparkes, et al.^62^. The constructs were transformed into *Agrobacterium tumefaciens* (AGL1) using electroporation, and grown at 28 degrees for 24-48 hours prior to being resuspended in the infiltration buffer (50 mM MES, 0.5% (w/v) glucose, 100 μM acetosyringone; pH 5.6) to a final OD_600_ of 0.1. The Agrobacterium was infiltrated into the upper leaves of 3-4 week old *N. benthamiania* plants which had been grown in a constant temperature room at 22°C with a 16-h light/8-h dark cycle. After 3-4 days the infiltrated leaves were examined using a Nikon A1Si confocal microscope (Nikon Plan Apo VC 60x NA1.2 water-immersion objective). The 488nm laser was used for excitation of GFP with the 521/50 nm band pass filter for fluorescence detection. The 561 nm laser was used for excitation of mCherry with the 595/50 nm band pass filter used to detect emitted fluorescence. Images were processed using the Nikon NIS elements software package.

### Transient *in planta* expression and confocal microscopy of rust effector candidates

Gene coding sequences without the predicted signal peptide sequence (SignalP 4.1) and stop codon were synthesised as single gBlock gene fragments (Integrated DNA Technologies, Coralville, Iowa, USA). Each gBlock included CACCATG at the 5’ end for cloning into the pENTR/D-TOPO Gateway entry vector (Invitrogen, Carlsbad, CA, USA) and to provide a start codon to initiate translation. The DNA was cloned directly into the entry vector, then transferred into pB7YWG2 containing the cauliflower mosaic virus (CaMV) 35S promoter, C-terminal EYFP gene and 35S terminator^63^ using LR Clonase II (Invitrogen). The final constructs were verified by DNA sequencing and then transferred by electroporation into the *Agrobacterium* strain GV3101 (pMP90). For *in planta* expression, *Agrobacterium* cultures were prepared to an optical density at 600 nm of 1.0 in 10 mM MES (pH 5.6) buffer with 10 mM MgCl_2_ and 200 μM acetosyringone, and then infiltrated into *N. tabacum* leaves. Infiltrated plants were kept in a 25°C growth room with a 16-hour day length. Tobacco leaf sections were imaged 2 days after agroinfiltration on a Zeiss confocal LSM 780 microscope using a 40× water immersion objective (LD C-Apochromat 40×/1.1W Korr M27) with excitation at 514 nm. EYFP fluorescence was detected at 520–600 nm and autofluorescence of the chloroplasts was detected at 605–720 nm or 650–720 nm. Zen 2012 digital imaging software (Zeiss, Oberkochen, Germany) was used during image acquisition.

### Immunoblot analysis

Two days after agroinfiltration, tobacco leaf tissue was frozen in liquid nitrogen and ground in 2x Laemmli buffer with 0.2 M dithiothreitol. The protein samples were boiled for 5 min, centrifuged to pellet leaf debris, then separated by sodium dodecylsulphate–polyacrylamide gel electrophoresis and transferred by electroblotting to a nitrocellulose membrane. The membrane was blocked in 5% skim milk then probed with anti-GFP (7.1 and 13.1; Roche, Penzberg, Germany), followed by sheep anti-mouse IgG antibody conjugated to horseradish peroxidase. Labelling was detected using the Amersham ECL Western Blotting Detection Reagent and the ImageQuant LAS 4000 CCD camera system (GE Healthcare Life Sciences, Little Chalfont, UK).

## Acknowledgements

We thank Sophien Kamoun and his group for useful discussions. We thank Brendan Kidd and Laura Davies for their valuable feedback. JS is supported by an OCE Postdoctoral Fellowship.

## Author contribution

J.S., P.D., K.S. and J.T. planned and designed the research. J.S. developed and implemented the machine learning method. A-M.C., K.D, B.P and D.G performed localization experiments. All authors analysed and interpreted the data and results and wrote the manuscript.

